# Discovery of potential drug-like compounds against Viral protein (VP40) of Marburg Virus using pharmacophoric based virtual screening from ZINC database

**DOI:** 10.1101/2021.05.13.444037

**Authors:** Sameer Quazi, Javed Malik, Komal Singh Suman, Arnaud Martino Capuzzo, Zeshan Haider

**Affiliations:** GenLab BioSolutions Private Limited, Bangalore, Karnataka, India; Department of Zoology, Guru Ghasidas Vishwavidyalaya, Bilaspur, Chhattisgarh, India; Department of Veterinary Sciences, University of Milan, Italy; Centre of Agricultural Biochemistry and Biotechnology (CABB), University of Agriculture Faisalabad, Pakistan

**Keywords:** MARV virus, VP40, Favipiravir, ZINC database, Virtual pharmacophore screening, Molecular docking, Drug design

## Abstract

Marburg virus (MARV) has been confirmed to cause extreme hemorrhagic fever (HFM) in human and animals. The effective and suitable vaccine to treat the MARV virus is not commercialized in public and demands rigorously tested on several scales. This research used a CADD (Computer-Aided Drug Design) computational based technology to find novel drug-like compounds that could inhibit the replicating of the VP40. The pharmacophoric features bases screening was done using an online computational based software, “ZINC Pharmer”. We retrieved about 32456 compounds mainly focused on the properties of pharmacophores from the ZINC database. Lipinski’s rule was also used to predict these drug-like compounds. as well as molecular coupling-dependent screening and selection of VP40 screening ligand complexes based on S-rank (lower than reference) and value of root, mean square (RMSD) (bottom) to examine for reference) using the Molecular Working Environment (MOE) machine. As a result, 100 compounds were found to have a close interaction with MARV VP40, followed by the Binding energy (BE) analysis of these 100 compounds. Only 50 were the strongest binding energy than favipiravir [reference inhibitor] after using the MOE-LigX algorithm to compare their binding energy. After that, ADMET analysis predicted only five compounds (ZINC95457352, ZINC38752258, ZINC38752253, ZINC39272175, and ZINC38752377) passed the ADMET parameters and have the strongest inhibitory effect against the MARV’s VP40. It has been suggested that these “drug-like” candidates have an increased ability to inhibit MARV replication, leading to treatment of MARVD.

## 1. INTRODUCTION

The Marburg virus disease [MARVD] is a human pandemic disease caused by MARV. The overall fatality percentage of MARVD varies from 23% to 90%, depending upon the epidemic condition of severity ^1^. MARV belongs to the family of Filoviridae, is an enveloped, nonsegmented, single-strand, and negative-strand RNA virus. The MARV has a small genome size, about 17kb to 19kb in length. The genome of MARV consisted of about seven different open reading frames (ORFs), like an additional viral protein, i.e., VP40, VP35, nucleoprotein, glycoprotein, and viral polymerase enzymes ^2,3^. The filoviridae family consisted of different types of viruses like ebolavirus, Marburgvirus, both of these viruses caused hemorrhagic fever and sometimes caused deadly illnesses in humans and animals. While other strains of filoviridae viruses like striavirus, cuevavirus, and thamnoviurs non-virulent to humans and animals ^4^.

The MARV filovirus was first identified after the epidemics in Germany and Serbia, respectively, in 1967 ^5^. After that, the MARV virus outbreak was discovered in different African regions. Later on, in 1999, an epidemic was a breakout in the democratic public of cango. This MARV outbreak brought out significant economic growth loss due to its highest 83% fatality rate in human communities ^6^. After five years, again, the epidemic of MARV was found in Angolia in 2005. It brought out a higher number of losses in Angolia due to the more effective strain of MARV. According to a survey, the fatality rate was more than 90% ^7^. Moreover, the impact of MARV’s outbreak still was not disappeared, while after some years, a big evaluation of MARV again appeared in different areas_of Uganda early in 2007. While in later 2008, some patients with MARV were also reported in the united states and Netherland. In short, some cases were also reported in 2012, 2013. 2017, 2019 in a diverse area of Uganda ^8^.

The favipiravir inhibitor has been earlier described as a potent inhibitor of MARV protein 40. Accordingly, the favipiravir inhibitor was selected to detect ligands from the ZINC database using the online pharmacophores-based platform ZINC Pharmer. Computer methods and virtual screening play a decisive starring role in drug development. High throughput virtual screening is a well-believed aspect of drug production ^9^. Innovative computational systems such as database pharmacophoric ligand screening, structure-based screening, molecular docking, and simulation dynamics could identify the novel drugs compounds that can be used as blocker agent against.

MARV’s VP40 catalytic active site. As a result of these computational biological tools, a modern study will be developed to discover combinations with close interaction, increased binding strength, and strong inhibitory effects with MARV VP40. These drug-like compounds are said to have a better potential for inhibiting MARV replication, leading to the development of treatment with MARVD.

## 2. MATERIALS AND METHODS

### 2.1 Computational based analysis

#### 2.1.1 Protein structure prediction, evaluation, and validation

The MARV VP40 protein (accession number APQ46231.1) sequence has been derived from the NCBI database. The three-dimensional structure was predicted using homology simulation after analysis of its primary sequence. The three-dimensional structure of the MARV VP40 protein was predicted by using MODELLER v9.25, a homology-based computer software (ul Qamar et al., 2020). The highest DOPE score [Discrete Optimized Protein Power] was used to select the premium structure. Besides, the PROCHECK analysis program was used to assess the accuracy of the projected 3D structure of the VP40.

### 2.2 Pharmacophore-based virtual screening of inhibitor of VP40

The steric and digital pharmacophore properties arrangement is crucial to ensure the best molecular interactions with the respective organic target and inhibit its natural function ^10^. The “ZINC-pharmer” software was used for the pharmacophore-based simulated scan of drug-like compounds from the ZINC online database ^11^. The pharmacophore ligand-receptor features using the three-dimensional (three-dimensional) configuration of MARV VP40 and the extension ratio of the favipiravir inhibitor “SDF” were automatically developed ZINC Pharmer ^12^. This screening type is essential for detecting potent target inhibitors, predicting new drug-like compounds, and promoting similar evaluation ^13^. The drug-like properties were by using Lipinski’s law. Under this law, drug-like properties must have a log P-value < five, a molecular weight < 500 Da, H-binding acceptors (HBA) and H-binding donors (HBD) < five. The general rule describes the molecular properties necessary for a drug’s pharmacokinetics in the human body ^14^. The connections’ main results were selected to be included in a new file to further analyze the interactions. The Molecular Operating Environment (MOE) program was used to build the three-dimensional test database ^15^.

### 2.3 Molecular docking

All the recovered compounds were docked to MARV VP40 to further test these drug-like compounds. MOE tools were used to perform ligand elimination, hydrogen insertion, and energy minimization of VP40 protein to reference molecular docking ^16^. The catalytic site of VP40 was discovered using the site search algorithm developed in the MOE package. The 2D interaction analysis algorithm was used to find and analyze the interaction between ligands and VP40. The molecular docking was performed between the ligands and VP40 by selecting different parameters (Rescoring function one and rescoring function two: London dG, location: triangle comparator, maintenance: 2, and refinement: field of pressure). The Favipiravir inhibitor S-score and root mean square deviation (RMSD) scores were used to select the novel compounds. The S-value calculates the evaluation effort that measures the ligand’s affinity for the receptor using the traditional MOE evaluation function. The RMSD method is used to compare the conformation of the coupled confirmation to the reference-related conformation. The Compounds that showed the lowest S and RMSD values were selected for further analysis ^17^. For further study, the binding energy of those selected compounds was also calculated using MOE software. The target protein receptor’s binding affinity with the ligand and the ligand’s hydrophobic and polar interactions with the target protein receptor are the main parameters to identify the small molecule’s practical effect on the target protein. Its 5 to 15 kcal/mol range represents a close contact between ligand and receptor ^18^. The compound with the same or greater binding affinity than the reference inhibitor was considered for further assessment of ingestion, release, metabolism, excretion, and toxicity profiles. The basic and biological properties of small drugs molecules have also been measured using the pkCSM server.

### 2.4 Validation and ADMET analysis

The pkCSM^19^ server was used to evaluate the ADMET (absorption, digestion, metabolism, excretion, and toxicity) properties of selected target compounds

## 3. Result and discussion

Several experimental trials have been conducted in the search for a MARV vaccine. However, there is currently no drug being created to successfully manage this pathogen ^4^. As a result, finding a low-cost antiviral medication that controls MARV is critical. A progressive approach to drug discovery is ensuring that novel drug-like compounds are effective against viral infections. Traditional drug research approaches are unreliable and time-consuming ^17^. Favipiravir, a broadspectrum antiviral drug, is currently used to treat MARV in vitro.

Additionally, in a computer simulation, the preferred Favipiravir inhibitor blocked the biological activity of VP40 by blocking its active online catalytic website ^20^. Despite the wider debate, the latest review centered on the screening of completely interactive and pharmacophores-based databases, molecular coupling, and drug similarity profiles. This is the most current technique for discovering active compounds that inhibit the proteolytic action of VP40 on the MARV catalytic pocket and can be used as medicines.

### 3.1 Structural prediction, evaluation, and validation of VP40

The viral protein of MARV VP40 consisted 303 number of amino acids. The physiochemical analysis revealed that the MW of VP40 was 106 kDa, while the GRAVY and instability index was −0.296 and 27.03, respectively. Furthermore, utilizing homology-based programmed modeler 9.25, the three-dimensional structure (3-D) of VP40 is expected. The total sequence similarity between the prototype “C5b0” and the template “C5b0” was about 93 percent. The best 3D structure was expected, and the Ramachandran plot assessment analysis showed that most of the amino acids (215 AA) are in the right place (94.3%). For further review, there’s no need to improve and format the analysis (figure 1).

**Figure 1.**
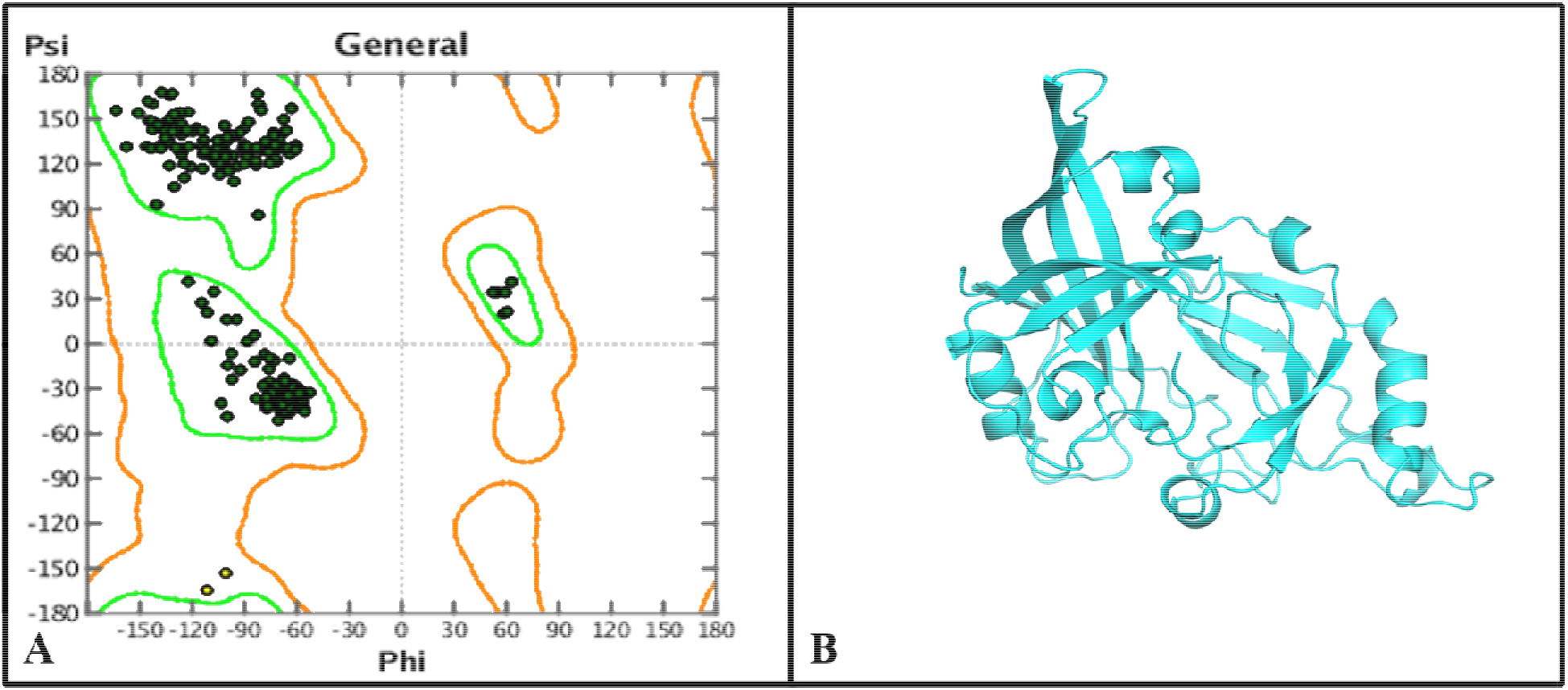
The 3D structure prediction of MARV VP40 (A) Ramachandran plot assessment shows the highest number of amino acids are in favourable regions (B) shows the graphical representation of 3D structural VP40.

### 3.2 Pharmacophore-based virtual screening and database preparation

Virtual screening is a fast and efficient way to identify new drug compounds in CADD. The ZINC pharmacophore software was used to screening the millions of drug-like combinations from the ZINC database. The pharmacophore complex model was created by the following parameters, i.e. hydrophobic (X= 4.85, Y= −0.20, Z= 0.00 and radius = 1.00), H acceptor (X= 2, Y= 0.16, Z= 0.00 and radius = 0.50), and H donor (X= 4.6, Y= −1.35, Z= 0.00 and radius = 0.50). The high quality with similar properties to the favipiravir inhibitor was obtained from the ZINC database (Figure 2). The overall 32456 compounds were retrieved from the ZINC database. Just 1266 drug-like compounds were chosen after implementing Lipinski’s rule of five. The minimization algorithm in the MOE programme was used to build a ready-to-dock catalogue.

**Figure 2.**
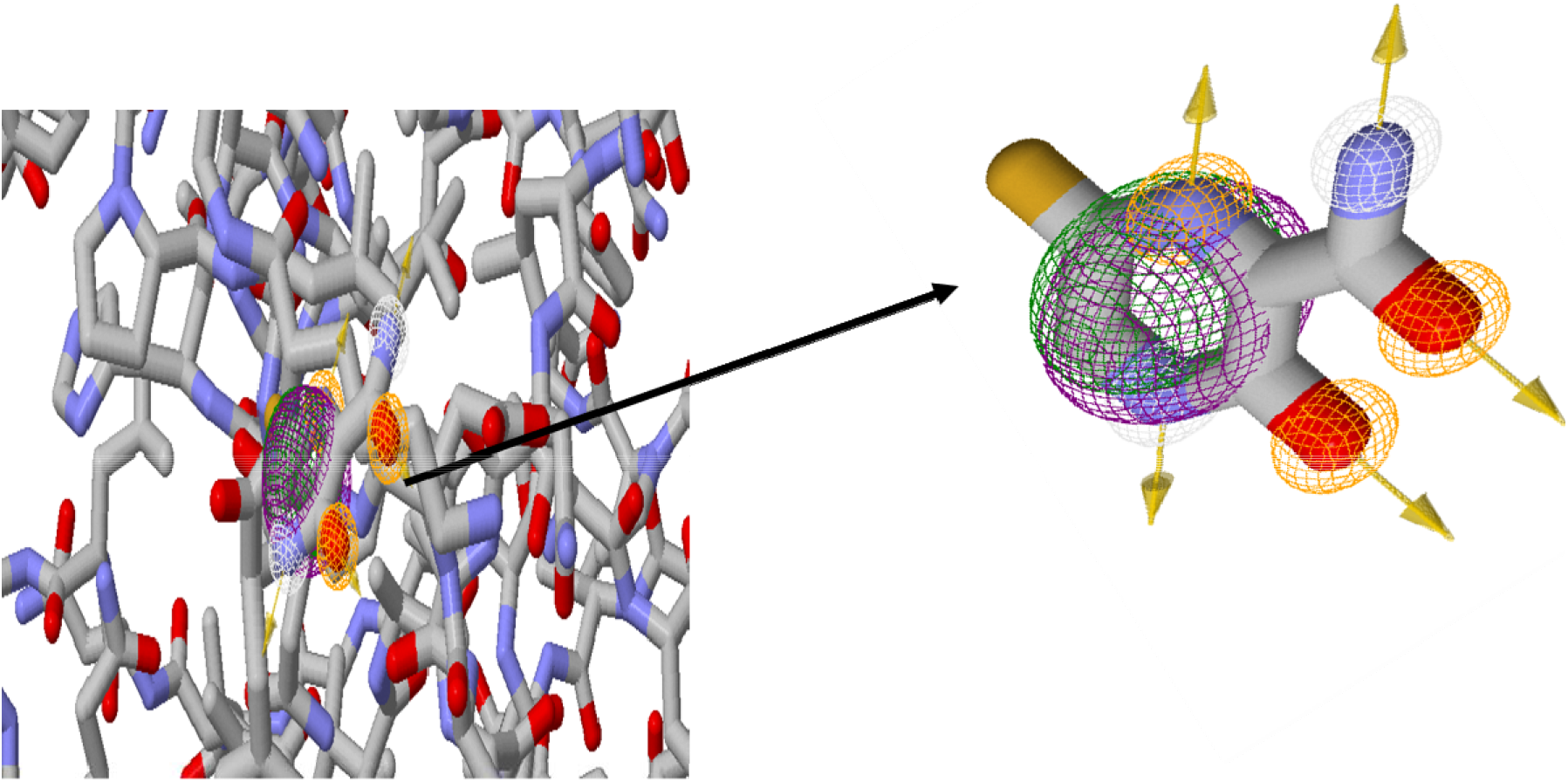
Graphical representation of pharmacophoric complex (VP40 + Favipiravir) generated by ZINC pharmer tool.

### 3.3 Molecular docking

It has been determined that molecular docking is essential in drug discovery. The active site finder algorithm determined the active site of MARV VP40 in the MOE package, which is the gateway to drug discovery. The VP40 protein was docked with by MOE, and the lowest S-score and RMSD docked complex ranked top of the list. The lowest S-score (−10.89) and RMSD (2.0) was selected for screening the novel drug-like small molecules. Furthermore, the library of druglike compounds was docked with VP40 protein. The uppermost 100 small molecules that have the lowest S-score and RMSD were selected for further study. The MOE Ligplot algorithm was used to calculate the binding ratio of these 100 compounds to VP40. The affected bonds’ bond strength was determined using the GB / VI Generalized Bond Volume Required (MOE) algorithm. As seen in Table 1, compounds with a stronger interaction with MARV VP40 than the reference inhibitor were chosen. The ADMET profile evaluation included fifty drugs like small molecules out of a total of 100.

**Table 1:** The Lipinski’s rule and molecular docking score of finally selected compounds.

| Docking Scores |  |  |  | Drug likeness scores |  |  |  |  |
| --- | --- | --- | --- | --- | --- | --- | --- | --- |
| ZINC-ID | S-value | RMSD | Binding Energy | MW | LogP | HBD | HBA | tPSA |
| ZINC95457352 | -13.50 | 1.99 | -17.73 | 252.3 | -1.74 | 4 | 4 | 94.85 |
| ZINC38752258 | -14.67 | 0.99 | -15.45 | 235.25 | 1.46 | 1 | 3 | 55.63 |
| ZINC38752253 | -14.50 | 1.12 | -14.30 | 136.13 | -1.57 | 2 | 2 | 55.43 |
| ZINC39272175 | -12.50 | 1.09 | -14.60 | 241.30 | 0.50 | 3 | 2 | 65.39 |
| ZINC38752377 | -12.35 | 1.79 | -17.23 | 245.33 | 1.14 | 3 | 2 | 65.39 |

### 3.4 Validation and ADMET analysis of hit compounds

The properties of ADMET were also analyzed on the fifty drug-like compounds using the pkCSM server. Only five small drug-like molecules passed the ADMET test. The Blood and Brain Barrier (BBB) is a difficult gift for the endothelial cells that keep the mind from taking drugs. The BBB is important in the field of drug development ^21^. Oral bioavailability is a significant factor in selecting active and effective drug against a specific patient ^22^. The following properties of ADMET analysis are very important for discovering effective drug compounds; therefore, this parameter has a significant role in the virtual screening of novel drugs like compounds against a specific disease or a pathogen.

These ADMET parameters such as P-glycoprotein substrate/inhibitor, Blood-brain barrier substrate/inhibitor (positive or negative), Human intestinal preparation (positive or negative), CaCo2 permeability (positive or negative), Renal Organic cation transporter (inhibitor/substrate), CYPs enzyme (inhibitor/non-inhibitor), and AMES toxicity (carcinogenic/non-carcinogenic) are the considered more significant parameters for the selection of good and effective drug-like compound. A good and effective drug must be passed the ADMET parameters. Out of fifty drugs like compounds best, five compounds were fulfilled the Admet requirements. These five (ZINC95457352, ZINC38752258, ZINC38752253, ZINC39272175, and ZINC38752377) druglike small molecules have significantly accepted the ADMET parameters (Table 2). The following parameters were used to select drug development compounds: ZINC ID, docking ranking, RMSD rating, binding energy value, VP40 binding residue, Lipinski’s results (Table 1), and ADMET study (Table 2). The compounds selected may be used as a novel, structurally distinct, and potentially active VP40 replication inhibitors (Figure 3).

**Figure 3.**
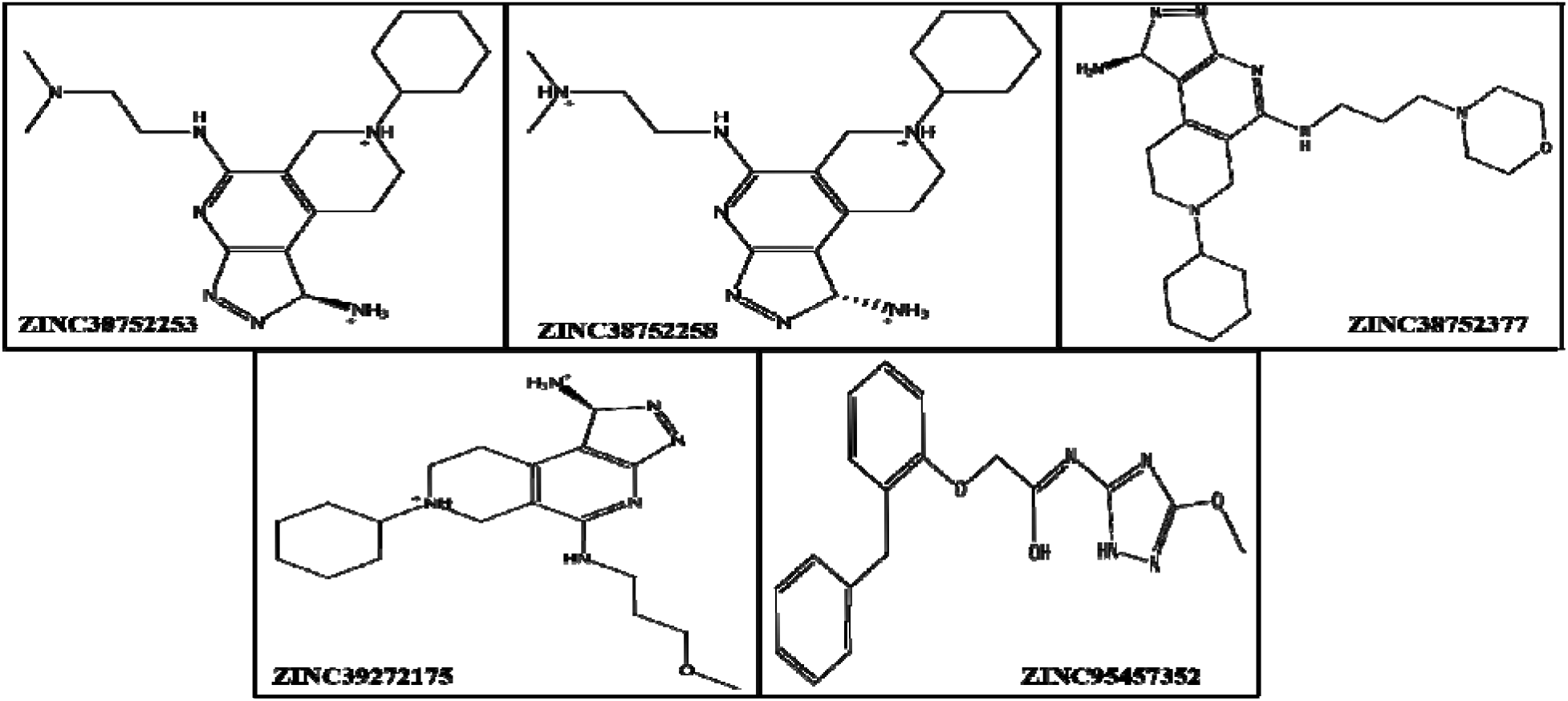
Representation of 2D structures of the novel drug-like compounds.

**Table 2:**
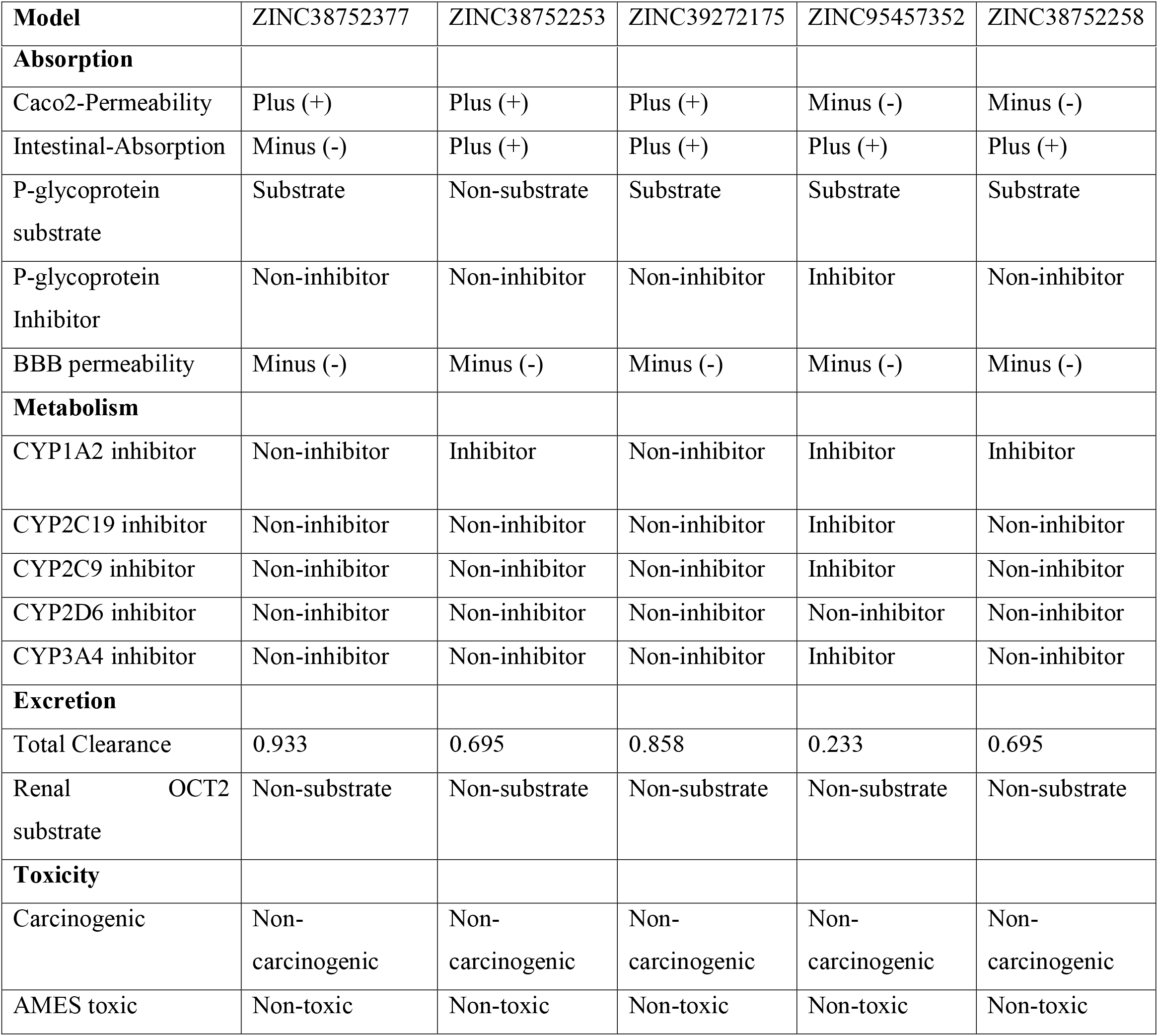
ADMET profiling of small drug-like molecules.

|  |  |  |  |  |  |
| --- | --- | --- | --- | --- | --- |
| **Model** | ZINC38752377 | ZINC38752253 | ZINC39272175 | ZINC95457352 | ZINC38752258 |
| **Absorption** |  |  |  |  |  |
| Caco2-Permeability | Plus (+) | Plus (+) | Plus (+) | Minus (-) | Minus (-) |
| Intestinal-Absorption | Minus (-) | Plus (+) | Plus (+) | Plus (+) | Plus (+) |
| P-glycoprotein substrate | Substrate | Non-substrate | Substrate | Substrate | Substrate |
| P-glycoprotein Inhibitor | Non-inhibitor | Non-inhibitor | Non-inhibitor | Inhibitor | Non-inhibitor |
| BBB permeability | Minus (-) | Minus (-) | Minus (-) | Minus (-) | Minus (-) |
| **Metabolism** |  |  |  |  |  |
| CYP1A2 inhibitor | Non-inhibitor | Inhibitor | Non-inhibitor | Inhibitor | Inhibitor |
| CYP2C19 inhibitor | Non-inhibitor | Non-inhibitor | Non-inhibitor | Inhibitor | Non-inhibitor |
| CYP2C9 inhibitor | Non-inhibitor | Non-inhibitor | Non-inhibitor | Inhibitor | Non-inhibitor |
| CYP2D6 inhibitor | Non-inhibitor | Non-inhibitor | Non-inhibitor | Non-inhibitor | Non-inhibitor |
| CYP3A4 inhibitor | Non-inhibitor | Non-inhibitor | Non-inhibitor | Inhibitor | Non-inhibitor |
| **Excretion** |  |  |  |  |  |
| Total Clearance | 0.933 | 0.695 | 0.858 | 0.233 | 0.695 |
| Renal OCT2 substrate | Non-substrate | Non-substrate | Non-substrate | Non-substrate | Non-substrate |
| **Toxicity** |  |  |  |  |  |
| Carcinogenic | Non-carcinogenic | Non-carcinogenic | Non-carcinogenic | Non-carcinogenic | Non-carcinogenic |
| AMES toxic | Non-toxic | Non-toxic | Non-toxic | Non-toxic | Non-toxic |

### 3.5 Receptor-ligands interaction analysis

The S-docking score determines the strength of the connection between the receptor amino acid and ligands atoms. Based on the docking score (S-value), binding energy, and other ADMET analysis, there are five small drugs like compounds selected as novel effective molecules. The five drugs’ molecules like ZINC95457352, ZINC38752258, ZINC38752253, ZINC39272175, and ZINC38752377, have represented a strong interaction complex with active amino acids of MARV’s VP40. The 3D docking conformation and proposed interaction mechanism of the respective ligand with VP40 are shown in Figure 4. The overall different amino acids like VAL-44, LEU-169, ILE-39, GLU-74, ALA-71, LYS-71, VAL-209, ALA-102, THR-99, and ALA-97 are the most active amino acid found that were involved in making hydrogen bonding and polar interaction with ligands.

**Figure 4.**
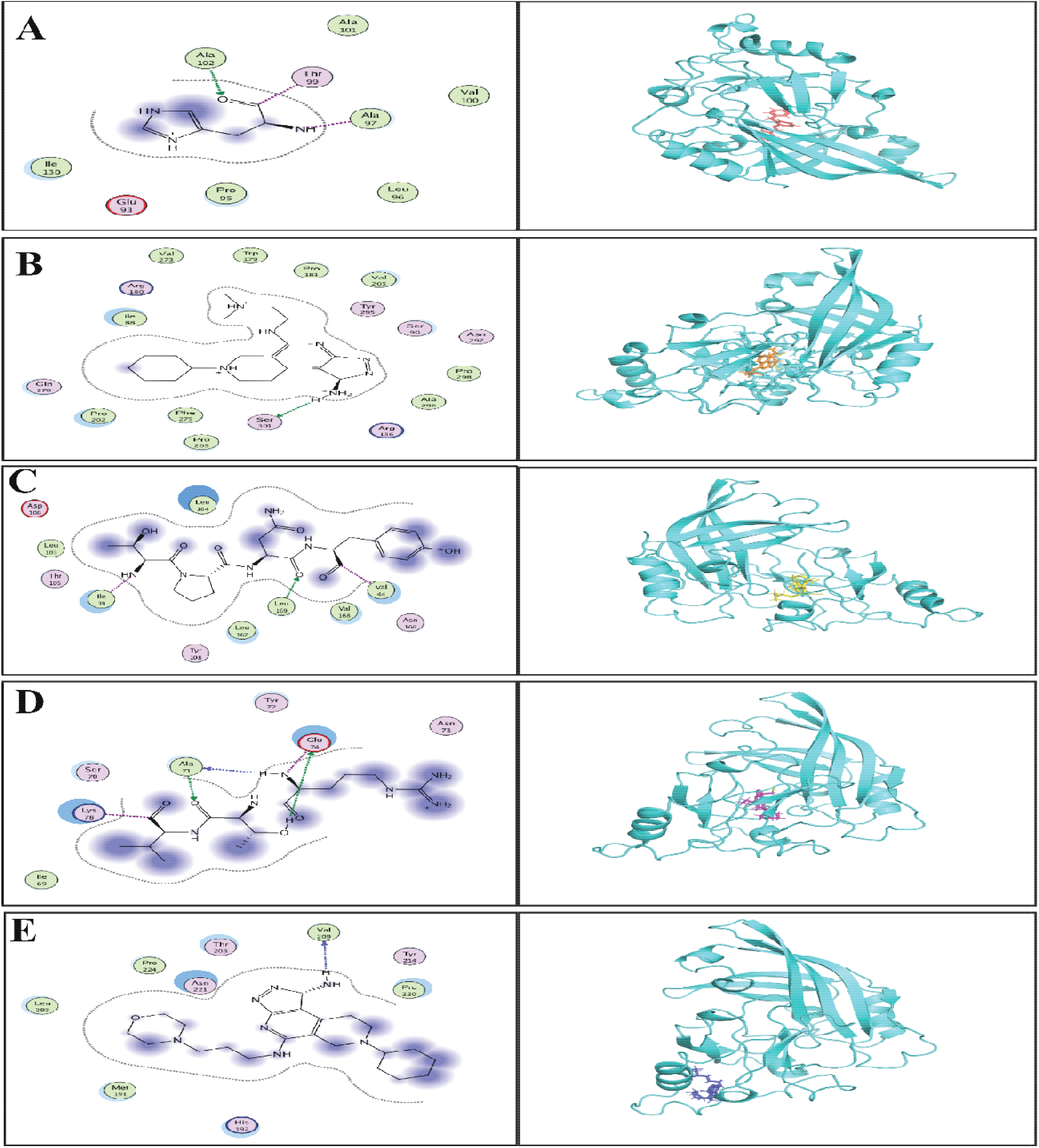
The docking confirmation of the best five drugs like compounds with the target VP40 protein of MARV. (A) represents ZINC95457352 docking to the VP40. The right side of the 2D conformation reveals that ALA-102, THR-99, and ALA-97 have heavy interaction with the ligand molecule, while the majority of the molecules are involved in binding surface area. Simultaneously, the 3D (left side) depicts the ligand conformation (red colour) in a receptor active bag (Cyan colour) (B) signifies the docking posed of ZINC38752258 against the VP40. The 2D conformation (right side) shows SER-301 is making strong interaction with ligand molecule, while the rest of the molecules involved in binding surface area. Simultaneously, the 3D (left side) shows the ligand conformation (orange colour) in an active pocket of the receptor (cyan colour) (C) describes ZINC38752253’s docking with VP40. The right side of the 2D conformation reveals that VAL-44, LEU-174, and ILE-39 have a strong interaction with the ligand molecule, while the majority of the molecules are involved in binding surface area. Simultaneously, the 3D (left side) portrays the ligand conformation (yellow colour) in a receptor active bag (cyan colour) (D) defines ZINC39272175’s docking with VP40. GLU-74, LYS-78, and ALA-71 are making heavy interactions with the ligand molecule in the 2D conformation (right side), while the majority of the molecules are involved in binding surface space. Simultaneously, the 3D (left side) depicts the ligand conformation (magenta colour) in a receptor active bag (cyan colour) (E) represents ZINC38752377’s docking with VP40. The right side of the 2D conformation shows VAL-209 having a close interaction with the ligand molecule, while the rest of the molecules are involved in binding surface space. Simultaneously, the 3D (left side) depicts the ligand conformation (blue colour) in a receptor active bag (cyan color).

## 4. CONCLUSION

Several experimental trials have been conducted in the search for a MARV vaccine. However, there is currently no drug or vaccine in progress to successfully manage this pathogen. As a result, finding a low-cost antiviral medication that controls MARV is critical. A progressive approach to drug discovery is ensuring that novel drug-like small molecules are successful against viral diseases.

Traditional drug research approaches are ineffective and time-consuming. Due to this reason, the present research’s primary goal was to use the ZINC database to conduct pharmacophore-based simulated scanning, molecular docking of selected compounds, and binding interaction testing against the MARV VP40. The five compounds ZINC95457352, ZINC38752258, ZINC38752253, ZINC39272175, and ZINC38752377, demonstrated a good interaction with the MARV VP40 active site. These findings suggest that these compounds could have the ability to be used as a MARV drug. It would be extremely useful in the research and design of in vivo drugs.

## CONFLICT OF INTERESTS

All the authors declared that there is no conflict of interest in the presented research paper.

## Notes

### Competing Interest Statement

The authors have declared no competing interest.

